# MTOR promotes basal cell carcinoma growth through atypical PKC

**DOI:** 10.1101/2020.08.24.264598

**Authors:** Rachel Y. Chow, Taylor M. Levee, Gurleen Kaur, Daniel P. Cedeno, Linda T. Doan, Scott X. Atwood

## Abstract

Advanced basal cell carcinomas (BCCs) are driven by the Hedgehog (HH) pathway and often possess inherent resistance to SMO inhibitors. Identifying and targeting pathways that bypass SMO could provide alternative treatments. Here, we use a combination of RNA-sequencing analysis of advanced BCC tumor-normal pairs and immunostaining of human and mouse BCC samples to identify an MTOR signature in BCC. Pharmacological inhibition of MTOR activity in BCC cells significantly reduces cell proliferation without affecting HH signaling. Similarly, treatment of *Ptch1*^*fl/fl*^; *Gli1*-*Cre*^*ERT2*^ mice with everolimus reduces tumor growth and aPKC activity, suggesting that MTOR promotes tumor growth by activating aPKC and demonstrating that suppressing MTOR could be a promising target for BCC patients.

## INTRODUCTION

Basal cell carcinoma (BCC) is the most common of cancers, with nearly 5 million new cases in the United States every year^[1]^. BCCs result from aberrant activation of the Hedgehog (HH) signaling pathway, an important pathway normally involved in embryonic development and adult tissue homeostasis^[2]^. Smoothened (SMO) inhibitors such as vismodegib are commonly used to suppress tumor growth in advanced cases. Unfortunately, SMO inhibitor treatment is only effective in ∼40% of advanced patients^[3]^, with ∼20% of patients who do respond eventually developing resistance each year^[4]^. Developing therapies to bypass SMO inhibitor resistance is a critical need and an active area of investigation, especially as inappropriate HH pathway activation also drives other cancers such as rhabdomyosarcoma, medulloblastoma, and basal cell carcinoma^[5,6,7]^.

Normally, HH ligand binds to the cholesterol transporter Patched1 (PTCH1), freeing up the G-protein coupled receptor SMO to inhibit SUFU and allowing for activation of the GLI transcription factors to translocate into the nucleus and activate target genes involved in proliferation, migration, and invasion^[2-8]^. Most patients who develop BCCs possess inactivating *PTCH1* (∼70%) or activating *SMO* (∼20%) mutations which drive uncontrolled HH signaling^[9]^, making SMO a natural target to treat a majority of cases. Resistance to SMO inhibitors primarily occurs through secondary mutations in SMO that either prevent drug binding or result in constitutive activation even when drug is bound^[10,11]^. Recent work on circumventing SMO inhibition has concentrated on shutting down GLI activation, where preclinical targeting of aPKC^[10,12]^, HDAC1 ^[13,14]^, MKL1^[15]^, and MEKK2/3^[16]^ have all shown some efficacy.

Targeting other GLI responsive signaling pathways is an area of growing interest, however, untangling their myriad interactions to define their mechanism of action is complex. For instance, loss of primary cilia in advanced BCCs can, in some cases, shut down HH signaling and concomitantly increase MAPK pathway activation, resulting in a switch from BCC to squamous cell carcinoma^[17]^. This mutual antagonism between RAS/MAPK and HH signaling can drive SMO inhibitor resistance and MAPK inhibitors can suppress tumor cell growth when the RAS/MAPK pathway is dominant^[18]^. MTOR is another major oncogenic player that has been associated with uncontrolled proliferation, resistance of cell death, evasion of immune destruction, and dysregulated cell metabolism^[19]^. Whether MTOR acts upstream, alongside, or downstream of the HH pathway is complicated by different results. In esophageal adenocarcinoma, MTOR functions through S6K1 to phosphorylate GLI1 and promote its transcriptional activity^[20]^. However, studies in neuroblastoma demonstrate that inhibition of the MTOR/S6K1 pathway suppresses cancer growth but does not affect GLI1 expression^[21]^. Additionally, in pancreatic and ovarian cancer cells, HH signaling has been shown to induce DYRK1B expression, which leads to activation of the MTOR/AKT pathway^[22]^.

Here, we provide evidence that an MTOR signature is significantly enriched in both human and mouse BCCs. We demonstrate that *in vitro* and *in vivo* inhibition of mTor results in significant reduction in BCC growth independent of aPKC-mediated activation of Hh signaling. Our results suggest that MTOR operates downstream of GLI1 and may be a viable target to treat advanced BCC patients.

## METHODS

### Ethics statements

Human clinical studies were approved by the Ethics Committee of the University of California Irvine. All human studies were performed in strict adherence to the Institutional Review Board (IRB) guidelines of the University of California Irvine.

### RNA Sequencing Analysis

RNA-seq data were obtained from patient-matched advanced human BCC patients^[10]^. RNA-Seq data were aligned as previously described^[10]^. The NCBI Reference Sequence (RefSeq) databases were used as reference annotations to calculate values of reads per kilobase of transcript per million mapped reads for known transcripts (RPKM). RPKM values were then log2-transformed and heat map analysis was used to visualize the differential gene expression. Pathway enrichment terms from RNA-seq data were obtained using Enrichr^[46]^.

### Human Samples

Written informed consent was obtained for all archived human samples and was reviewed by the University of California Irvine IRB. Human normal epidermis and BCC samples were collected from UC Irvine Medical Center. Paraffinized samples were sectioned with a rotary microtome (Leica RM2155) at 7 μm for analysis. Samples were deparaffinized as described by Abcam and antigen retrieval was performed using a Tris-EDTA buffer (10 nM Tris base, 1 mM EDTA, 0.05% Tween 20, pH 9.0) at 60°C overnight.

### Cell Culture

ASZ001 cells were grown in 154CF media containing chelated 2% FBS, 1% Penicillin-streptomycin, and 0.07mM CaCl2 (Life Technologies).

### Hedgehog assay

ASZ001 cells were plated to confluence, serum-starved (SS), and treated with either DMSO or varying concentrations of Rapamycin (0.1 nM, 1 nM, 10 nM, and 100 nM) (Fisher Scientific), OSI-027 (5 μM, 10 μM, 25 μM, and 100 μM) (Fisher Scientific), and Everolimus (2 nM, 10 nM, 50 nM, and 250 nM) (Fisher Scientific) for 24 hours. RNA was isolated using the Direct-zol RNA MiniPrep Plus (ZYMO Research). Quantitative RT-PCR was performed using the iTaq Univer SYBR Green 1-Step Kit (Bio-Rad) on a StepOnePlus Real-time PCR system (Applied BioSystem). The fold change in *Gli1* mRNA expression (forward: 5′-GCAGGTGTGAGGCCAGGTAGTGACGATG-3′, reverse: 5′-CGCGGGCAGCACTGAGGACTTGTC-3′) was measured using ΔΔCt analysis with *Gapdh* (forward: 5′-AATGAATACGGCTACAGCAACAGGGTG-3′, reverse: 5′-AATTGTGAGGGAGATGCTCAGTGTTGGG-3′) as an internal control. Experiments were repeated three times and ran in triplicates.

### Growth assay

ASZ001 cells were seeded at 2000 cells/well into 96-well plates. After 48 hours, cells were treated with DMSO or varying concentrations of Rapamycin, OSI-027, and Everolimus (refer to HH assay) for 2, 4, and 6 days. Growth assay was performed with MTT (Sigma-Aldrich) per manufacturer’s protocol. Plates were analyzed with a Bio-Tek uQuant MQX200 plate reader. Experiments were repeated three times in 6 wells each.

### Mice

All mice were housed under standard conditions and animal care was in compliance with the protocols approved by the Institutional Animal Care and Use Committee (IACUC) at University of California Irvine. *Ptch1*^*fl/fl*^; *Gli1*-*Cre*^*ERT2*^ mice were administered 100uL of 10mg/mL tamoxifen (Sigma) intraperitoneally for 3 consecutive days at 6 weeks of age. 5 weeks later, mice were treated with either DMSO or Everolimus (3mg/kg) intraperitoneally for 7 consecutive days. The final volume of all injections was 100 μL. At the end of treatment, mice were sacrificed and their back skin collected, fixed in 4% paraformaldehyde (PFA) for 30 minutes at room temperature, washed with PBS, immersed in 30% sucrose at 4°C overnight, and frozen in optimal cutting temperature (OCT) compound (Sakura Finetek). Samples were then cryo-sectioned (CryoStar NX50) at 14 μm for analyses. Five mice were used for each treatment condition.

### Micro-tumor assessment

Mouse sections were H&E stained per standardized protocol and images were taken at 200x magnification on an AmScope microscope with an AmScope MU500B digital camera. Tumor size was measured using FIJI software. Micro-tumors were assessed in mouse back skin as total tumor size per square area. More than 50 tumors were measured in each of 5 mice.

### Immunofluorescence staining

Skin sections were blocked using 10% BSA and 0.1% Triton X-100 in PBS for 1 hr at room temperature. The following antibodies were used: mTor (rabbit, Cell Signaling Technology 2983S, 1:400), Gli1 (rabbit, Santa Cruz Biotechnology sc-20687, 1:500), Gli1 p-T304 (rabbit, 1:200) ^[34]^, Krt14 (chicken, Fisher Scientific 50-103-0174, 1:5000), aPKC (rabbit, Santa Cruz Biotechnology sc-216, 1:500), aPKC p-T410 (rabbit, Santa Cruz Biotechnology sc-12894-R, 1:200), and aPKC p-T560 (rabbit, Abcam ab62372, 1:300). Sections were mounted with ProLong Diamond AntiFade Mountant with DAPI (Invitrogen). Images were acquired on a Zeiss LSM700 confocal microscope with a 63x oil immersion objective. FIJI was used to determine the average pixel intensity over five distinct tumors within a given skin section. Images were arranged with FIJI and Adobe Illustrator.

### Statistics

Statistical analyses were done using two-tailed *t* test or two-way ANOVA using GraphPad Prism.

## RESULTS

### MTOR pathway expression is significantly enriched in advanced human BCCs

To identify alternative pathways that drive BCC growth, we reanalyzed our bulk-level RNA sequencing data from 14 tumor-normal pairs of advanced BCC patients^[10]^. 1602 genes were upregulated by two-fold or more when differential gene expression was averaged across all 14 samples **(Figure 1A; Supplementary Table 1)**. KEGG analysis of upregulated genes indicated the expected cancer-related terms such as cell cycle, DNA replication, cell metabolism, genes involved in BCC, and HH signaling **(Figure 1B; Supplementary Table 2)**. Interestingly, the MTOR pathway showed significant enrichment with the related PI3K-AKT and HIF-1 pathways also present **(Figure 1B; Supplementary Table 2)**. MTOR-related pathways were even more prominent with Kinase Enrichment Analysis^[23]^ of the upregulated genes where MAPK, AKT, GSK3B, PLK1 and S6K kinase terms all significantly enriched, along with the expected CDKs **(Figure 1C; Supplementary Table 2)**. When we analyzed gene expression of the MTOR pathway, MTORC1 complex components and downstream targets showed significant upregulation such as AKT1S1, S6K1, and EIF4EBP1, whereas MTOR gene expression itself was significantly increased in only a subset of tumors **(Figure 1D; Supplementary Table 3)**.

**Figure 1.**
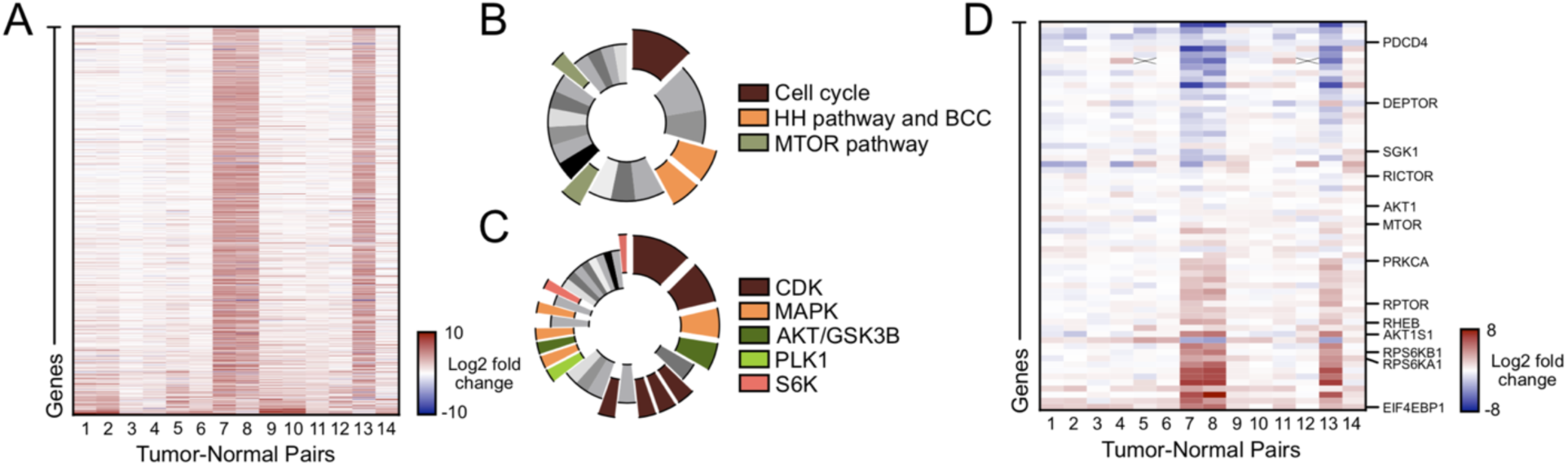
The MTOR pathway is differentially expressed in advanced human BCCs. **A)** Heat map of differentially expressed genes upregulated by 2-fold or more in advanced human BCCs compared to patient-matched normal skin. **B)** KEGG analysis of differentially expressed genes showing significant terms as indicated. **C)** Kinase Enrichment Analysis of differentially expressed genes showing significant kinases as indicated. **D)** Heat map of differentially expressed mTOR pathway components in advanced human BCCs compared to patient-matched normal skin. Select genes denoted on the right. X, no data.

### MTOR is upregulated in human and mouse BCCs

To validate MTOR expression at the protein level in BCC tumors, we immunostained both human and mouse BCC tumor samples and compared them to normal epidermis. Human BCC tumors showed significant enrichment of MTOR in nodular human BCC tumors compared to normal epidermis **(Figure 2A-B)**. However, individual tumor immunostaining showed a large variation in protein expression with some tumors not showing enrichment compared to normal epidermis, a similar pattern to the RNA-seq data analysis **(Figure 1D)**. To analyze mTor expression in mice, we utilized a *Ptch1*^*fl/f*l^; *Gli1*-*Cre*^*ERT2*^ mouse model in which BCC tumors predominantly arise from *Gli1*-positive regions within the hair follicle bulge and secondary hair germ^[24]^. BCC tumors were allowed to grow for five weeks post-tamoxifen treatment and formed predominantly from the hair follicle regions. mTor immunostaining showed significantly increased expression in tumors compared to either the normal epithelium or normal hair follicle **(Figure 2C-D)**. A similar variation in staining was observed in both mouse and human tumors. Collectively, these results indicate that the MTOR pathway is overexpressed in both human and mouse BCCs.

**Figure 2.**
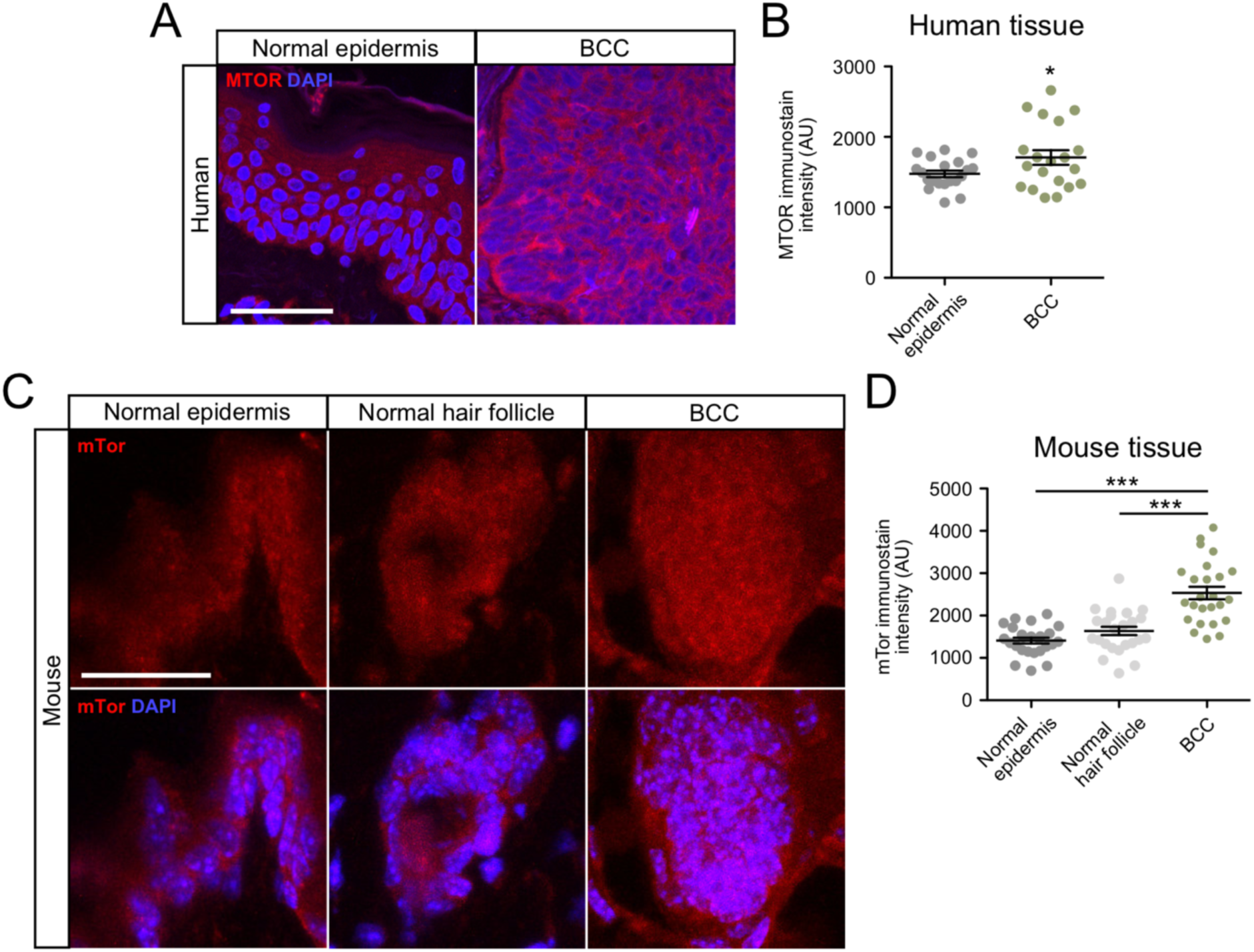
MTOR is overexpressed in human and mouse BCC. **A)** Immunofluorescent staining of indicated markers in human normal epidermis and nodular BCC. Scale bar, 50 μm. **B)** Quantification of MTOR immunostain intensity (n = 5 different points of measurement from 4 individual samples). AU, arbitrary units. **C)** Immunofluorescent staining of indicated markers in normal epidermis, normal hair follicle, or BCC derived from *Ptch1*^*fl/fl*^; *Gli1*-*Cre*^*ERT2*^ mice. Scale bar, 25 μm. **D)** Quantification of mTor immunostain (n = 5 different points of measurement from 5 mice). Error bars represent SEM. Significance was determined by unpaired two-tailed *t* test. *, p < 0.05; **, p < 0.01; ***, p < 0.001.

### mTor inhibition suppresses murine BCC cell growth but not Hh signaling

We next wanted to assay whether mTor inhibition suppresses Hh signaling and tumor cell growth. We treated ASZ001 mouse BCC cells with three different mTor inhibitors that are in various stages of clinical use: rapamycin, OSI-027, and everolimus. Rapamycin and everolimus act as allosteric inhibitors, while OSI-027 acts as a competitive ATP inhibitor^[25, 26, 27]^. Rapamycin and OSI-027 treatments did not result in significant changes in Hh signaling as assayed by *Gli1* mRNA expression, whereas everolimus treatment resulted in a slight but not significant increase in *Gli1* expression **(Figure 3A)**. These data complement previous reports that show mTor expression is Hh-dependent in ASZ001 mouse BCC cells^[28]^, suggesting that mTor operates downstream of the Hh pathway in BCC. Despite not significantly influencing Hh signaling, treatment with all three mTor inhibitors resulted in a dose-dependent decrease in BCC cell growth over time **(Figure 3B-D)**. Reduced proliferation of BCC cells undergoing rapamycin and everolimus treatment can be seen as early as two days after initial drug exposure, whereas OSI-027 treatment required at least four days to see a significant effect on tumor cell growth compared to DMSO vehicle control. Together, these data demonstrate that mTor inhibition can suppress BCC cell growth without altering *Gli1* expression, suggesting that mTor operates downstream of the Hh pathway.

**Figure 3.**
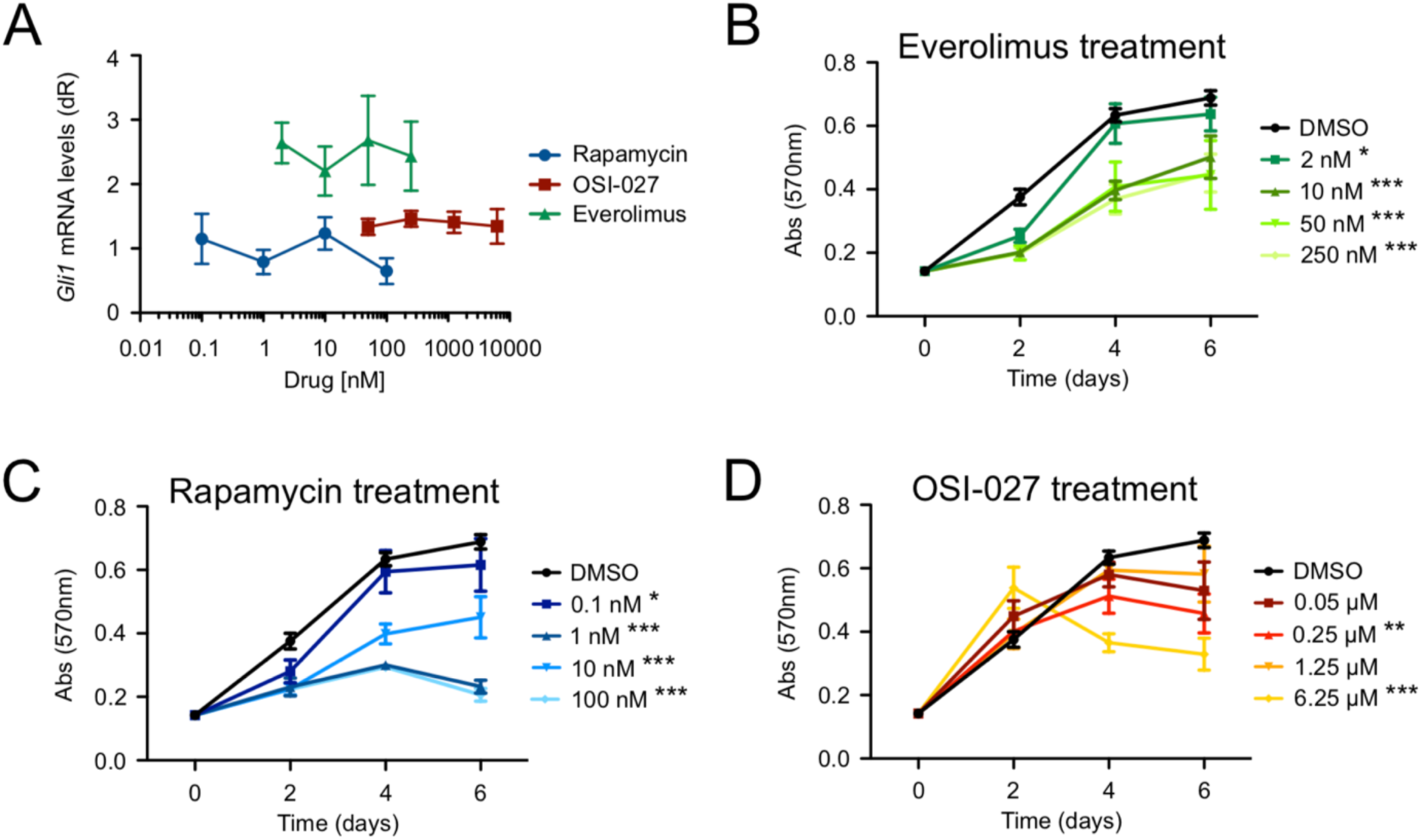
mTor inhibition suppresses BCC cell growth but not HH signaling. **A)** Gli1 mRNA levels of ASZ001 cells treated with DMSO or varying concentrations of Rapamycin, OSI-027, or Everolimus (n = 3 experiments). dR, delta reporter signal normalized to passive reference dye. **B-D)** MTT assay of ASZ001 cells treated with **B)** Everolimus, **C)** Rapamycin, or **D)** OSI-027 (n = 3 experiments). Abs, absorbance. Error bars represent SEM. Significance was determined by two-way ANOVA test. *, p < 0.05; **, p < 0.01; ***, p < 0.001.

### mTor function independently of aPKC to suppress murine BCC tumors

To explore whether mTor inhibition can be used as an effective BCC therapeutic, we grew BCC tumors in *Ptch1*^*fl/fl*^; *Gli1*-*Cre*^*ERT2*^ mice for 5 weeks after tamoxifen injection and intraperitoneally injected either DMSO or 3 mg/kg everolimus once a day for seven days. We used everolimus despite a slight increase in *Gli1* expression in ASZ001 mouse BCC cells because it is FDA approved and has been shown to be more effective in treating certain cancers and other diseases than rapamycin^[29, 30, 31, 32]^. Histological staining of the dorsal skin of everolimus-treated mice showed a significant reduction in total tumor area compared to DMSO controls **(Figure 4A-B)**. Gli1 immunostains showed no significant change in expression **(Figure 4C-D)**, corresponding to the quantitative PCR data from mouse BCC cells and reinforcing mTor’s function downstream of the Hh pathway.

**Figure 4.**
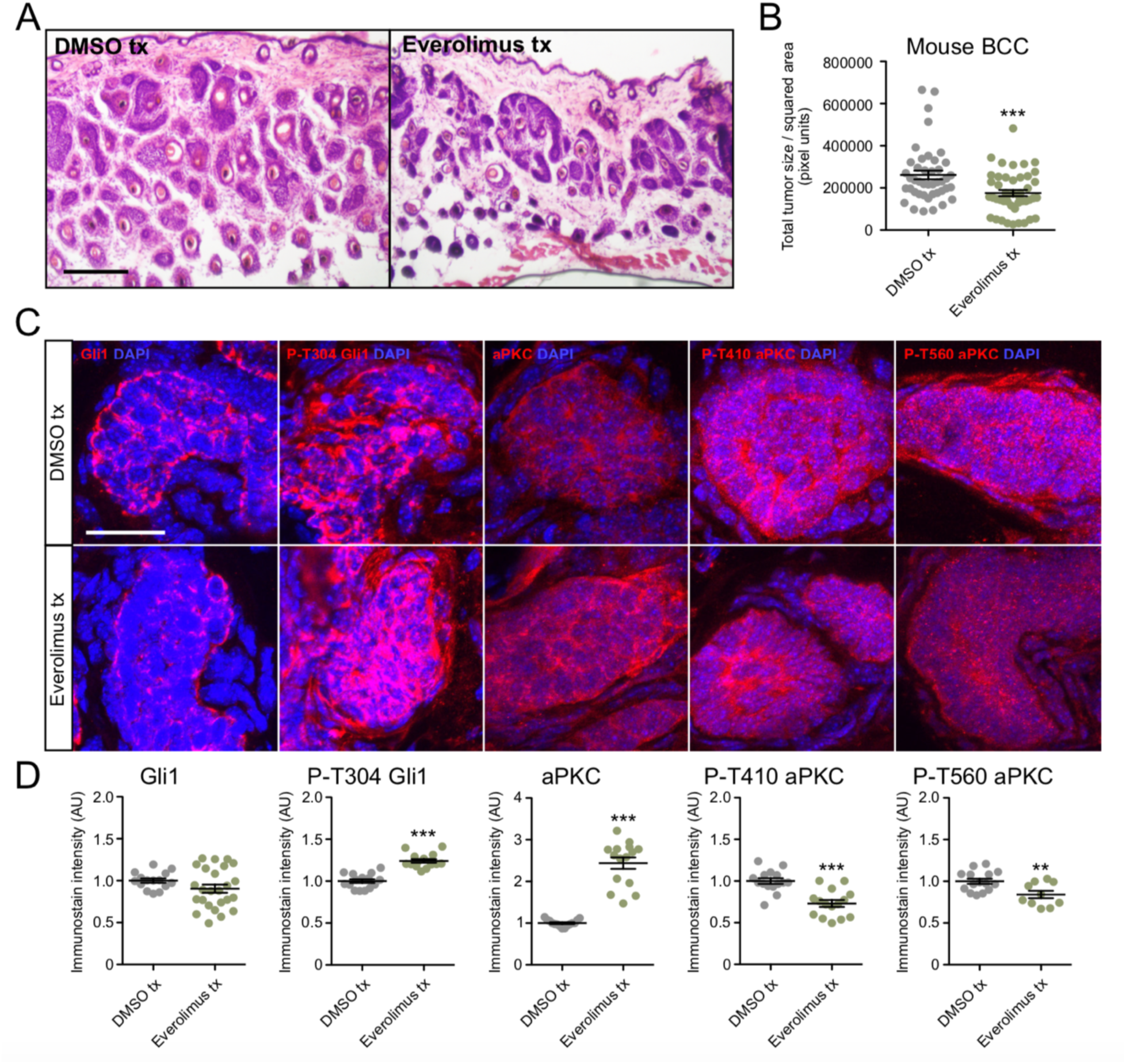
mTor inhibition suppresses murine BCC growth and aPKC activity. **A)** Hematoxylin and eosin staining of dorsal back skin collected from DMSO- or Everolimus-treated *Ptch1*^*fl/fl*^; *Gli1*-*Cre*^*ERT2*^ mice. Scale bar, 50 μm. **B)** Quantification of total tumor size per square area (n > 250 tumors from 5 mice). tx, treatment. **C)** Immunofluorescent staining of DMSO- or Everolimus-treated *Ptch1*^*fl/fl*^; *Gli1*-*Cre*^*ERT2*^ mice for the indicated markers. Scale bar, 25 μm. **D)** Quantification of immunostains (n = 5 tumors from 3 mice). AU, arbitrary units. Error bars represent SEM. Significance was determined by unpaired two-tailed *t* test. *, p < 0.05; **, p < 0.01; ***, p < 0.001.

To further delineate mTor’s mode of action, we assayed that status of aPKC, a Gli1 kinase that is necessary for high sustained Gli1 activity^[12]^. mTor has been shown to phosphorylate and activate aPKC at residue T560 in mouse embryonic fibroblasts^[33]^, while aPKC phosphorylates and activates Gli1 at residue T304 ^[12, 34]^. We observe reduced p-T560 aPKC immunostaining in everolimus-treated mouse BCC tumors, suggesting that mTor activity is important to phosphorylate this residue in BCC **(Figure 4C-D)**. We also observe a concomitant reduction in p-T410 aPKC, an activation site that is thought to be phosphorylated by Pdpk1^[35]^. However, total aPKC and p-T304 Gli1 expression were significantly enriched in everolimus-treated tumors **(Figure 4C-D)**, suggesting that mTor is likely promoting tumor growth independent of Gli1 and potentially through another aPKC target.

## DISCUSSION

How MTOR interacts with the HH pathway is unclear given reports that it can operate upstream, downstream, or parallel to the pathway in various cancers. For instance, in glioblastoma multiforme, MTOR inactivates GSK3β to prevent GLI2 ubiquitination, thereby promoting GLI2 stability and nuclear translocation^[36]^. This is likely not the case in BCC as we show MTOR inhibition does not significantly alter GLI1 expression. Our findings are more consistent with models where MTOR acts downstream of the HH pathway, such as in *Ptch1*^*+/-*^/*SKH-1* BCCs where Hh signaling promotes Sox9 expression to enhance mTor activity and tumor growth^[28]^. In fact, *SOX9* is significantly enriched in our bulk-level RNA sequencing data of advanced BCC patients **(Supplementary Table 1)**.

Our data and others suggest that MTOR acts downstream of the HH pathway to promote tumor growth, but MTOR’s mechanism of action is less clear. MTOR may converge on Cyclin D1 (CCND1) to directly promote BCC cell growth, which is also a target of the HH pathway^[37]^, as MTOR inhibition disrupts CCND1/CDK2 complexes^[38]^. Another possibility is that mTOR affects BCC growth via AKT1, an MTOR target that functions downstream of HH signaling in BCCs^[39, 40]^. MTOR phosphorylates AKT1 at S473^[38]^, and ASZ001 mouse BCC cells treated with intraconazole, a SMO inhibitor, reduces p-S473 AKT1 expression. Our data suggests MTOR phosphorylates and activates aPKC in BCC, but does not alter phosphorylation, suggesting that another aPKC target may be responsible for continued tumor growth^[41]^.

Cancer is heterogeneous and BCCs are no exception^[9, 10, 11]^. Our bioinformatics analysis and subsequent immunostaining suggests not all tumors possess an MTOR profile, where a subset of tumors showing strong upregulation while others displaying a more modest MTOR signature. This is not surprising as other signaling pathways are known to regulate BCC in conjunction with HH signaling, such as the WNT^[42]^, NOTCH^[43,44]^, and Hippo pathways^[45]^. As such, combination therapy may be an important step going forward to therapeutically treat advanced BCC patients.

## ACKNOWLEDGEMENTS

We would like to thank Sunny Wong for the *Ptch1*^*fl/fl*^; *Gli1*-*Cre*^*ERT2*^ mice. The work is funded by NIH grant R01CA237563 (SXA) and ACS Research Scholar Award RSG-19-089-01-DDC (SXA). The authors wish to acknowledge the support of the Chao Family Comprehensive Cancer Center Optical Biology Core Shared Resource, supported by the National Cancer Institute of the NIH under award number P30CA062203. The content is solely the responsibility of the authors and does not necessarily represent the official views of the NIH.

## AUTHOR CONTRIBUTIONS

S.X.A. and R.Y.C. conceived the project; S.X.A. supervised research; R.Y.C. performed experiments; T.L., G.K. and D.P.C. quantified and immunostained tumor data; L.T.D. collected and annotated human clinical samples. S.X.A. and R.Y.C. wrote the manuscript. All authors analyzed and discussed the results and commented on the manuscript.

---

**Supplementary Table 1. List of upregulated genes from RNA sequencing data of 14 tumor-normal pairs of advanced BCC patients**. List of genes corresponding to Figure 1A.

**Supplementary Table 2. KEGG and KEA analysis of upregulated genes from RNA sequencing data of 14 tumor-normal pairs of advanced BCC patients**. List of significant terms corresponding to Figure 1B.

**Supplementary Table 3. MTOR pathway gene expression from RNA sequencing data of 14 tumor-normal pairs of advanced BCC patients**. List of genes corresponding to Figure 1C.

